# A pathogenic DYT-THAP1 dystonia mutation causes hypomyelination and loss of YY1 binding

**DOI:** 10.1101/2021.08.03.454996

**Authors:** Dhananjay Yellajoshyula, Abigail E. Rogers, Audrey J. Kim, Sumin Kim, Samuel S. Pappas, William T. Dauer

**Affiliations:** Department of Neurology, University of Michigan, Ann Arbor, MI, USA; Molecular Cellular and Developmental Biology, University of Michigan, Ann Arbor, MI, USA; Cell and Molecular Biology, University of Michigan, Ann Arbor, MI, USA; Peter O’Donnell Jr. Brain Institute, University of Texas Southwestern Medical Center, Dallas, Texas, USA; Department of Neurology, University of Texas Southwestern Medical Center, Dallas, Texas, USA; Department of Neuroscience, University of Texas Southwestern Medical Center, Dallas, Texas, USA

## Abstract

Dystonia is a disabling disease that manifests as prolonged involuntary twisting movements. DYT-THAP1 is an inherited form of isolated dystonia caused by mutations in *THAP1* encoding the transcription factor THAP1. The phe81leu (F81L) missense mutation is representative of a category of poorly understood mutations that do not occur on residues critical for DNA binding. Here, we demonstrate that the F81L mutation (THAP1^F81L^) impairs THAP1 transcriptional activity and disrupts CNS myelination. Strikingly, THAP1^F81L^ exhibits normal DNA binding but causes a significantly reduced DNA binding of YY1, its transcriptional partner that also has an established role in oligodendrocyte lineage progression. Our results suggest a model of molecular pathogenesis whereby THAP1^F81L^ normally binds DNA but is unable to efficiently organize an active transcription complex.

## Introduction

DYT-THAP1 or DYT6 is an inherited form of isolated dystonia resulting from dominantly inherited mutations in *THAP1* (1). THAP1 is a ubiquitously expressed member of the THAP protein family of transcription factors that share an atypical zinc-dependent DNA-binding region, termed the “THAP” domain (2-4). THAP1 was first described as having a role in regulating cell-cycle and apoptotic genes in endothelial (HUVEC) cells (3, 5). Subsequent to the discovery of *THAP1* mutations in DYT6 dystonia (1), considerable work has focused upon identifying the THAP1 function and its regulated transcriptome in CNS tissue. This work demonstrates an essential role for THAP1 in CNS development and function (4, 6-9). THAP1 plays a pronounced role in CNS myelination via a cell-autonomous role in the development of the oligodendrocyte progenitor cells (OPC) into mature myelinating oligodendrocytes (OLs) (4). In addition to its role in OLs, THAP1’s role in normal neuronal function has been explored. THAP1 loss-of-function models demonstrate dysregulated striatal eIF2α signaling (6), abnormal cerebellar physiology (9) and elevated extracellular striatal acetylcholine (7). These studies also demonstrate that the THAP1-regulated transcriptome is highly tissue-dependent and likely dependent on additional transcriptional partners. The findings of Aguilo et.al emphasize this point, demonstrating that there is only a ∼10% overlap between THAP1-bound genes (ChIPseq) in mouse embryonic stem (mES) cells and genes differentially regulated from *Thap1* null mES (RNAseq)(10). THAP1 partners are therefore very likely to comprise a key regulatory element of THAP1 transcriptional activity in the CNS.

More than 100 putative *THAP1* mutations have been reported; most (73/113) are missense mutations (11), while fewer are indel mutations or cause early truncation (11-13). Many missense mutations occur in the N-terminal THAP domain, but relatively few are predicted to impact DNA binding directly. Studies that combine structure-function analyses and NMR studies demonstrate that DNA binding is mediated directly by 8 invariant residues (C5, C10, C54, H57, P26, W36, F58, and P78) and 5 additional residues (K24, R29, R42, F45 and T48) (3, 14, 15). Most of these residues reside on the DNA binding surface formed by the anti-parallel two-stranded β-sheet and the loop-helix-loop structure in the THAP domain (15).

Like all missense mutations that do not impact DNA binding residues, the mechanism whereby F81L impacts THAP1 function is poorly understood. Here, we demonstrate that THAP1^F81L^ exhibits impaired transcriptional activity. Similar to THAP1 null mice, THAP1^F81L^ mutant mice exhibit deficits in CNS myelination and abnormalities of compact myelin. Strikingly, the F81L mutation does not disrupt the DNA binding of THAP1. Rather, loss-of-function characteristic of THAP1^F81L^ is caused by significantly reduced binding of its transcriptional partner YY1, and corresponding decreases in the epigenetic marker H3K9ac at its target loci. These observations support a model whereby THAP1^F81L^ is normally bound to DNA but unable to organize an active transcription complex.

## Results

### The F81L DYT6 mutation disrupts THAP1 transcriptional activity

In prior work we generated and characterized a knock-in mouse line containing a floxed *Thap1*^*F81L*^ allele (4). The F81 residue does not bind DNA, but is part of the invariantly conserved AVPTIF motif in the THAP domain (Figure 1A-B) (15). These prior findings demonstrate that germline *Thap1*^*-/-*^ mice exhibit embryonic lethality (4, 10). In contrast, *Thap1*^*F81L/-*^ mice are born at expected Mendelian ratios and do not exhibit significant abnormalities of growth or lifespan (4). Considered together, these observations demonstrate that the *Thap1*^*F81L*^ allele complements *Thap1* function, but do not indicate whether F81L is a gain- or loss-of-function mutation.

**Figure 1.**
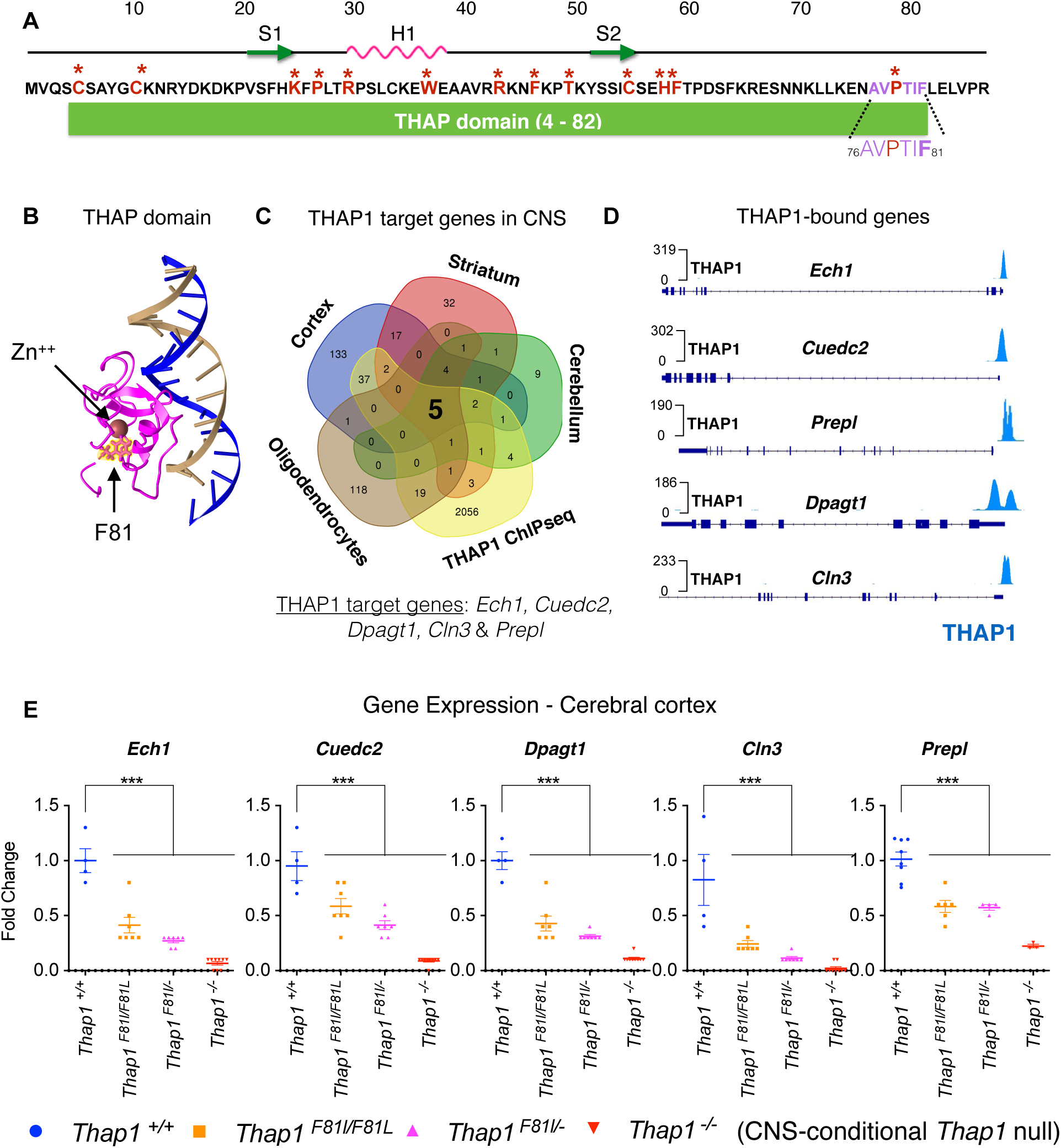
The DYT6 F81L mutation impairs THAP1 transcriptional activity. (A) Residues of the N-terminal THAP domain (residues 4-82). Residues involved directly in DNA binding are highlighted in red and starred. Also shown is the AVPTIF motif (highlighted in purple in inset) and the phe81 residue that is a focus of this study (highlighted in bold purple in the inset). Also illustrated is the location of the phe81 residue relative to the location of the DNA binding surface formed by the two β-strands (S1 and S2) and the α-helix motif (H1) forming the loop-helix-loop structure (15). (B) 3D model of the solution structure (15) of the complex between the THAP zinc finger of hTHAP1 and its specific *Rrm1* DNA target highlighting the location of the phe81 residue (yellow) generated using Cn3D (https://www.ncbi.nlm.nih.gov/Structure/CN3D/cn3d.shtml). (C) Venn diagram depicting the identification of target genes that are bound (ChIPseq) and regulated (*Thap1*^*+*/+^ vs *Thap1*^*-*/-^ differentially expressed genes (DEG) in CNS (cortex, striatum and cerebellum and OL lineage) by THAP1. The intersection of all datasets identified five THAP1-target genes *Ech1, Cuedc2, Dpagt1, Prepl* and *Cln3*. (D) Genome browser track (Integrative Genomics Viewer; https://igv.org) demonstrating THAP1 enrichment at the promoter region of its target loci *Ech1, Cuedc2, Dpagt1, Prepl and Cln3* (ENCODE dataset; K652 cells). (E) Significant reduction in the mRNA expression of *Ech1, Cuedc2, Cln3 and Dpagt1* in *Thap1*^*F81L/F81L*^, *Thap1*^*F81L/-*^ and N-CKO CNS (cerebral cortex) relative to control (*Thap1*^*+/+*^) as measured by qRT-PCR. mRNA expression for each individual gene was normalized to *Rpl19* expression and represented in the bar graph (mean ± SEM) as fold change (y-axis) for all genotypes (x-axis) with respect to *Thap1*^*+/+*^. One-way ANOVA for *Ech1* = F_(3,23)_ = 49.75, p<0.0001; *Cuedc2* = F_(3,23)_ = 35.55, p<0.0001; *Dpagt1* = F_(3,23)_ = 58.763, p<0.0001; *Cln3* = F_(3,23)_ = 20.53, p<0.0001 and *Prepl* = F_(3,23)_ = 28.13, p<0.0001; Dunnett’s multiple comparisons test: adjusted p value < 0.0001.

To further assess the effect of the F81L mutation, we sought to identify genes that are highly transcriptionally sensitive to THAP1. We examined 4 datasets that include genes significantly altered by THAP1 loss in brain tissue (separate datasets for cortex, striatum, and cerebellum), and in primary cultures of oligodendrocyte progenitor cells (16), as well as ChIP-seq data from ENCODE (Figure 1C and Table S1). We identified 5 genes (*Ech1, Cuedc2, Dpagt1, Prepl and Cln3*) that were common to all of these datasets (Figure 1C). THAP1 binds to the promoter of each of these genes (Figure 1D), and the expression of these genes is reduced more than 10-fold in the cerebral cortex of CNS-conditional *Thap1* null mice (*Thap1*^*flx/-*;^ Nestin-Cre or “N-CKO”; N-CKO vs *Thap1*^*+/+*^ ; one-way ANOVA; p < 0.0001 for all genes; N = 4; Figure 1E). The sensitivity of these genes to THAP1 in multiple contexts identified them as ideal candidates to explore the transcriptional impact of pathogenic *Thap1* mutations.

We examined the effect of the F81L mutation on the transcription of these 5 target genes (*Ech1, Cuedc2, Dpagt1, Cln3 and Prepl)*. We assayed the expression of these genes in cortical tissue from an allelic series of *Thap1 mutant mice* (*Thap1*^*+/+*^, *Thap1*^*F81L/F81L*^, *Thap1*^*F81L/-*^, *and Thap1*^*-/-*^). Expression of all five genes was significantly decreased in both *Thap1*^*F81L/F81L*^ and *Thap1*^*F81L/-*^ compared to *Thap1*^*+/+*^ cortical tissue (*Thap1*^*F81L/F81L*^ : > 2 fold decrease for all genes; N = 4; one-way ANOVA: p < 0.001 for *Ech1, Dpagt1, Prepl* and *Cln3 and p < 0*.*0016 for Cuedc2; Thap1*^*F81L/-*^ : > 5 fold decrease for *Ech1, Dpagt1* and *Cln3;* one-way *ANOVA: p < 0*.*0001* and *> 2 fold* decrease for *Cuedc2* and *Prepl*; N = 4; one-way ANOVA : p < 0.0001; Figure 1E). Expression of all 5 genes was further reduced by THAP1 deletion in the CNS (N-CKO) (Figure 1E). These results demonstrate that the F81L mutation significantly reduces THAP1 transcriptional activity, despite having no effect THAP1 expression (Figure S1). There were no significant differences in the expression of 4 (of 5) of these genes between *Thap1*^*F81L/F81L*^, *Thap1*^*F81L/-*^ cortical tissue, indicating that the F81 mutation has a marked impact on the transcription of these targets. These data, together with the viability of mice containing a single F81L allele, identify *Thap1*^*F81L*^ as a hypomorphic allele.

### The F81L DYT6 mutation causes CNS hypomyelination

We reported that conditional deletion of *Thap1* either from the CNS (nestin-Cre; N-CKO) or from oligodendrocytes (Olig2-Cre; “O-CKO”) delays myelination, overtly visible as hypomyelination in the juvenile CNS (4). We therefore explored whether the F81L loss-of-function mutation similarly causes myelination defects. We assessed the density of myelinated axons and myelin ultrastructure in the genu of the corpus callosum (CC) from P21 *Thap1*^*+/+*^, *Thap1*^*F81L/F81L*^, *Thap1*^*F81L/-*^ and N-CKO mice using transmission electron microscopy (TEM). Mice carrying the *Thap1*^*F81L*^ allele exhibited significant reductions in the density of myelinated axons in the CC (Figure 2A,D; 25% decrease in *Thap1*^*F81L/F81L*^ ; one-way ANOVA; p = 0.007 and 35% decrease in *Thap1*^*F81L/-*^ ; one-way ANOVA; p = 0.0018 and 70% decrease in N-CKO; one-way ANOVA; p < 0.0001). The observed hypomyelination did not result from axonal degeneration, as total axonal density (myelinated and non-myelinated combined) did not differ significantly between control, F81L and N-CKO genotypes (Figure 2E). The compact myelin formed also exhibited structural defects in DYT6 mutant CNS. The myelin sheaths of N-CKO, *Thap1*^*F81L/-*^ & *Thap1*^*F81L/F81L*^ mice were significantly thinner (larger g-ratio)(Figure 2B-C, F; 15% increase in g-ratio for both *Thap1*^*F81L/F81L*^ ; and *Thap1*^*F81L/-*^; one-way ANOVA; p < 0.0001 and 25% increase in g-ratio for N-CKO; one-way ANOVA; p < 0.0001). Loss of THAP1 function also increased the average diameter of myelinated axons, suggesting preferential loss in the myelination of smaller caliber axons (Figure 2B,G; ∼20% increase in the caliber of axons myelinated in *Thap1*^*F81L/ F81L*^, *Thap1*^*F81L/-*^ and N-CKO tissue ; one-way ANOVA; p < 0.0001).

**Figure 2.**
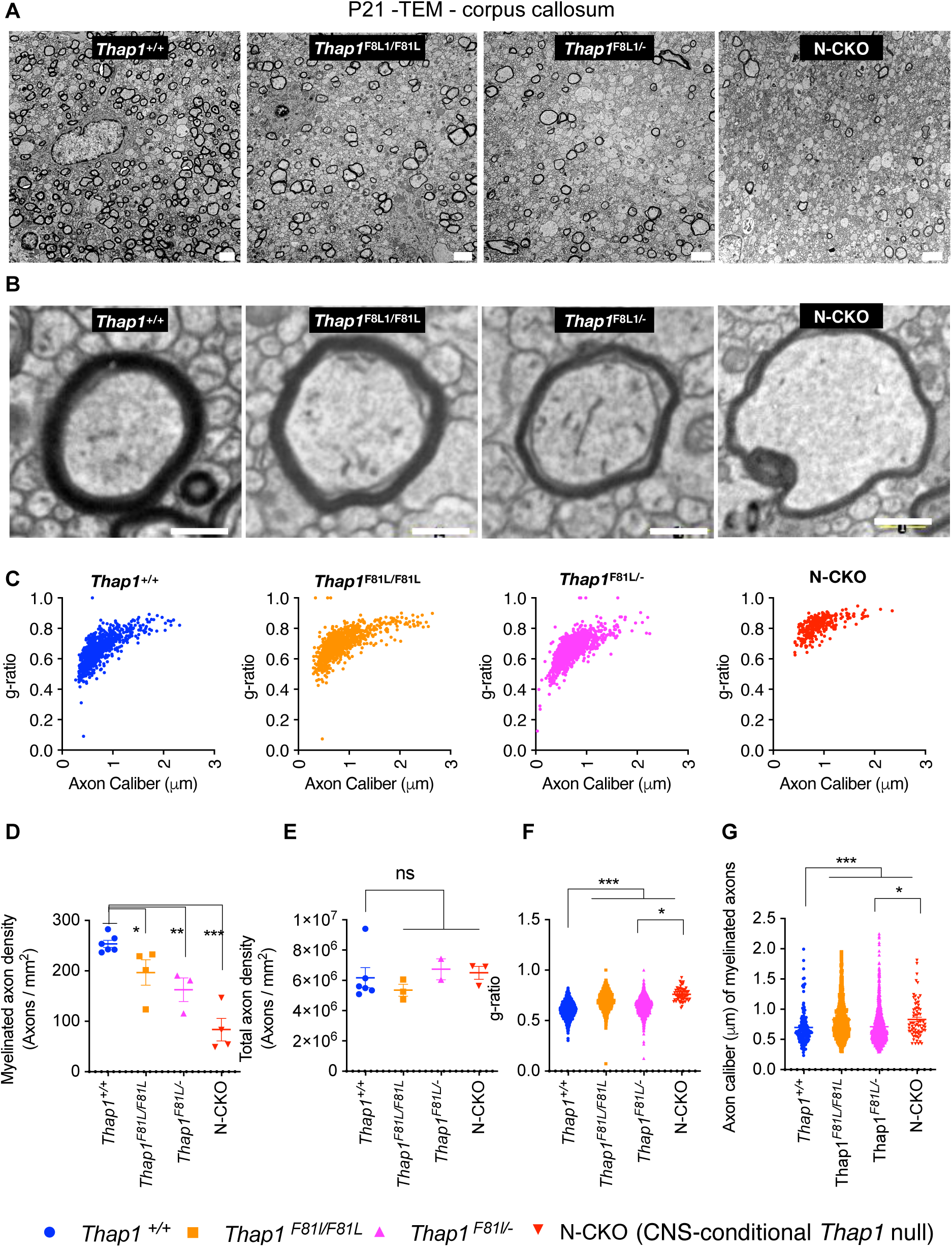
The DYT6 F81L mutation causes hypomyelination *in vivo*. (A-B) Representative transmission EM images at low (scale bar 4 μm; (A) and high magnification (scale bar 200 nm; (B) from the genu of corpus callosum (CC) at P21 from control (*Thap1*^*+/+*^), DYT6 homozygous (*Thap1*^*F81L/F81L*^), DYT6 hemizygous (*Thap1*^*F81L/-*^) and N-CKO (*Thap1*^*F81L*/-^; Nestin-*Cre*^*+*^) micr. (C) Graph depicting g-ratio (y-axis) represented in relation to axon caliber (μm; x-axis) for *Thap1*^*+/+*^, *Thap1*^*F81L/F81L*^, *Thap1*^*F81L/-*^ and *Thap1*^*F81L*/-^; Nestin-*Cre*^*+*^. (D) Quantification of the density of myelinated axons (y-axis; number of axons / mm^2^) represented as mean ± SEM for all genotypes (x-axis). *Thap1*^*+/+*^ = 253.8 ± 7.11, *Thap1*^*F81L/F81L*^ = 196.8 ± 26.33; *Thap1*^*F81L*/-^ = 162.6 ± 23.6; N-CKO = 83.5 ± 22.39; One-way ANOVA F_(3,13)_ = 16.73, p<0.0001, Dunnett’s multiple comparisons test: adjusted p value < 0.0001. (E) Quantification of the density of total axons (y-axis; number of axons / mm^2^) represented as mean ± SEM for all genotypes (x-axis). *Thap1*^*+/+*^ = 6.18 × 10^6^ ± 6.6 × 10^5^, *Thap1*^*F81L/F81L*^ = 5.35 × 10^6^ ± 3.86 × 10^5^; *Thap1*^*F81L*/-^ = 6.73 × 10^6^ ± 6.87 × 10^5^; N-CKO = 6.50 × 10^6^ ± 4.31 × 10^5^; One-way ANOVA F_(3,10)_ = 16.73, p=0.619, Dunnett’s multiple comparisons test: adjusted p value = 0.7183. (F) g-ratio represented in relation to genotype (x-axis) as mean ± SEM. *Thap1*^*+/+*^ = 0.60 ± 0.002, *Thap1*^*F81L/F81L*^ = 0.698 ± 0.0028; *Thap1*^*F81L*/-^ = 0.652 ± 23.6; N-CKO = 0.76 ± 0.007 One-way ANOVA F_(3,3035)_ = 205.1, p<0.0001, Dunnett’s multiple comparisons test: adjusted p value < 0.0001. (G) Axon caliber of myelinated axons (y-axis; μm of individual axons) represented as mean ± SEM for all genotypes (x-axis). *Thap1*^*+/+*^ = 0.69 μm ± 0.025, *Thap1*^*F81L/ F81L*^ = 0.828 μm ± 0.010; *Thap1*^*F81L*/-^ = 0.71 μm ± 0.008; N-CKO = 0.83 μm ± 0.03. One-way ANOVA F_(3,3035)_ = 205.1, p<0.0001, Dunnett’s multiple comparisons test: adjusted p value < 0.0001.

### The F81L mutation does not disrupt endogenous THAP1 binding at target loci

Having established that the F81L mutation impairs THAP1 function both transcriptionally and functionally at the level of myelination, we next explored the molecular mechanism responsible for this effect. Available evidence from structural analyses (3, 14, 15) (Figure 1B) indicates that the F81 residue does not participate directly in DNA binding. We assessed whether the mutation may nevertheless affect, perhaps indirectly, DNA occupancy of THAP1. Using quantitative chromatin immunoprecipitation (qChIP), we compared the binding of THAP1 and THAP1^F81L^ at several target loci. We performed qChIP analyses in clonal neural stem cell (NSC) lines because they are a uniform cell source common to the neural lineage. NSC were derived from *Thap1*^*+/+*^, *Thap1*^*F81L/-*^, *and Thap1*^*-/-*^ (*Thap1*^*flx/-;*^ *Nestin-Cre*) mouse lines. *Prior to qChIP, we assessed whether the transcription of our established target genes remained sensitive to THAP1 loss in these cells. Of the five THAP1 target genes, four* (*Ech1, Cuedc2, Dpagt1, Prepl)* exhibited significantly decreased expression in *Thap1*^*-/-*^ NSC cells (> 10 fold decreased expression for *Ech1* and *Cuedc2* and > 5 fold decreased expression for *Dpagt1* and *Prepl* in *Thap1*^*-/-*^ vs *Thap1*^*+/+*^; ANOVA: p < 0.0001 for *Ech1* and *Cuedc2;* Figure 3A) confirming that THAP1 is required for the expression of these genes in NSC. The other core THAP1 target gene, *Cln3*, showed negligible expression in NSC so was not further studied (data not shown). qChIP analyses confirmed that THAP1 is bound to the promoter region of *Ech1, Cuedc2* and *Prepl* loci in the NSC lineage, which was absent in THAP1 null NSC (Figure 3B). We detected very low amounts of THAP1 (∼ 10 fold lower) bound to the promoter of *Dpagt1* loci in NSC, leading us to focus on *Ech1, Cuedc2* and *Prepl* for further mechanistic studies in NSC. Specificity of qCHIP was further confirmed by lack of signal from the promoter of the active housekeeping gene *Gapdh* (Figure 3B) and lack of binding by an isotype specific negative control antibody (Goat IgG) to the target loci (Figure 3B). We next measured endogenous THAP1^F81L^ binding at *Ech1, Cuedc2* and *Prepl* loci in *Thap1*^*F81L/-*^ NSC and compared these data to THAP1 binding in *Thap1*^*+/+*^ NSC. In striking contrast to chromatin derived from *Thap1*^*-/-*^ NSC, F81L-THAP1 binding in *Thap1*^*F81L/-*^ NSC is comparable to that of *Thap1*^*+/+*^ cultures at the promoter regions of *Ech1, Cuedc2* and *Prepl* loci (Figure 3C), whereas no binding was detected at the control *Gapdh* loci (Figure 3C). Thus, the F81L DYT6 mutation does not significantly disrupt THAP1 DNA binding.

**Figure 3.**
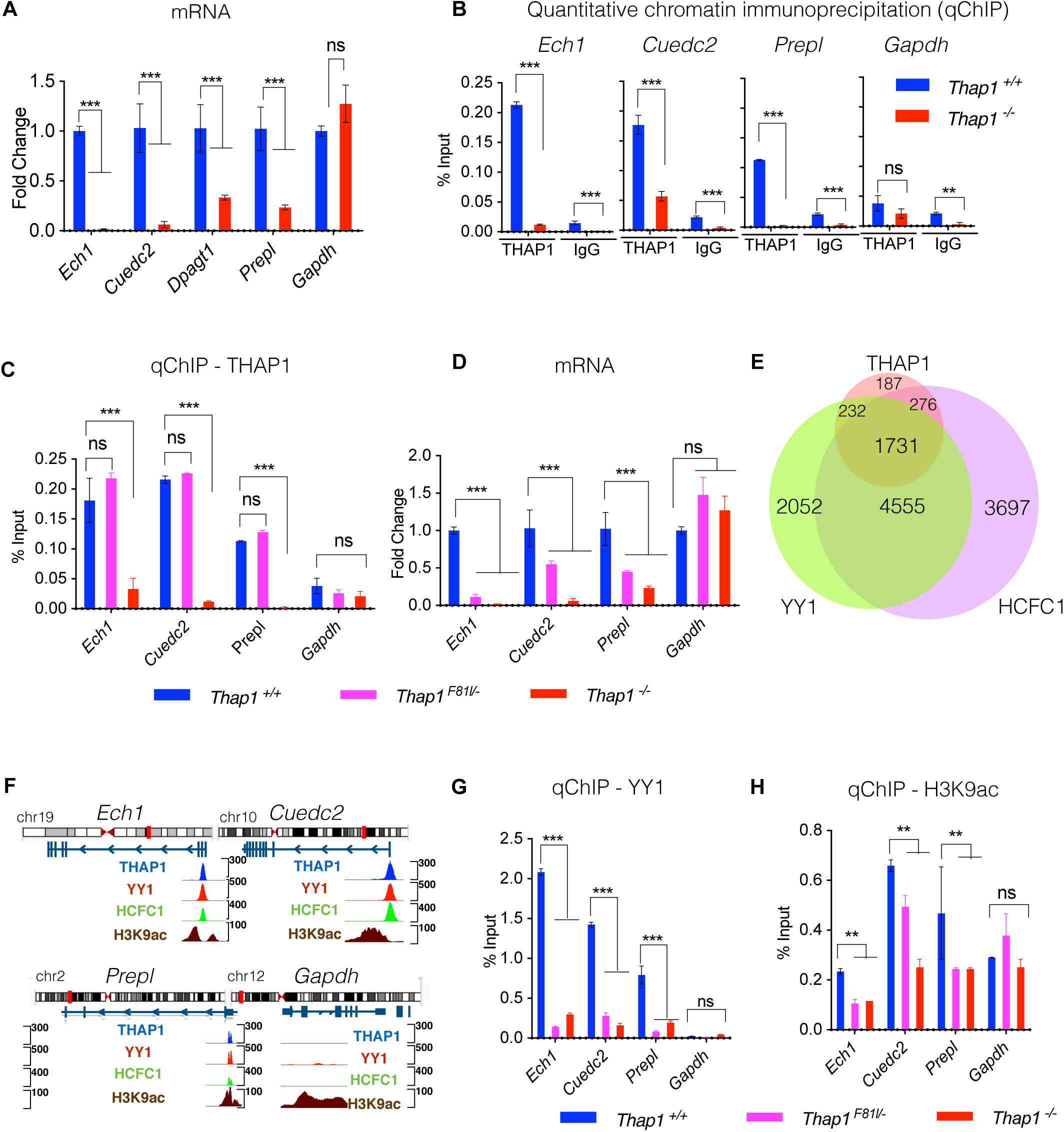
The DYT6 F81L mutation disrupts endogenous YY1 binding. (A) Significant reduction in the mRNA expression of *Ech1, Cuedc2, Dpagt1* and *Prepl* in *Thap1*^*-/-*^ NSC relative to *Thap1*^*+/+*^ NSC as measured by qRT-PCR (*Gapdh expression was unchanged)*. mRNA expression for each individual gene (x-axis) was normalized to *Rpl19* expression and represented in the bar graph (mean ± SEM) as fold change (y-axis) for *Thap1*^*-/-*^ relative to *Thap1*^*+/+*^. t-test; p<0.0001 for *Ech1, Cuedc2, Dpagt1* and *Prepl*; p = 0.37 for *Gapdh*. (B) Quantitative ChIP (qChIP) demonstrating the binding of THAP1 and isotype matched IgG antibody (x-axis) at the promoter region of target ge*nes* in NSC. Binding, represented as % input (y-axis), is demonstrated for THAP1 and normalized anti-goat IgG for chromatin isolated from *Thap1*^*+/+*^ and *Thap1*^*-*/-^ NSC cells for *Ech1, Ceudc2, Prepl and Gapdh* loci; ANOVA; p<0.0001 for *Ech1* and *Cuedc2* and p = 0.342 for *Gapdh*. (C) Quantitative ChIP (qChIP) demonstrating the binding of THAP1 at the promoter region of *Ech1, Cuedc2, Prepl and Gapdh* loci in NSC. Binding, represented as % input (y-axis), is demonstrated for THAP1 for chromatin isolated from *Thap1*^*+/+*^, *Thap1*^*F81L/-*^ and *Thap1*^*-*/-^ NSC cells for the promoter region of *Ech1, Cuedc2, Prepl and Gapdh* loci (x-axis). One-way ANOVA for THAP1 qChIP -*Ech1* - F_(2,6)_ = 18,65, p = 0.0027; *Cuedc2* - F_(2,6)_ = 1354, p value < 0.0001 and *Gapdh* - F_(2,6)_ = 11.72, p = 0.038; Dunnett’s multiple comparisons test: adjusted p value < 0.0001. (D) Significant reduction in the mRNA expression of *Ech1, Cuedc2* and *Prepl* but not *Gapdh* in *Thap1*^*F81L/-*^ and *Thap1*^*-/-*^ NSC relative to *Thap1*^*+/+*^ NSC as measured by qRT-PCR. mRNA expression for each individual gene (x-axis) was normalized to *Rpl19* expression and represented in the bar graph (mean ± SEM) as fold change (y-axis) for all genotypes with respect to *Thap1*^*+/+*^. One-way ANOVA for *Ech1* - F_(3,23)_, p<0.0001; *Cuedc2* - F_(3,23)_ = 22.36, p<0.0001 and *Gapdh* - F_(3,23)_ = 22.44 p = 0.0016; Dunnett’s multiple comparisons test: adjusted p value < 0.0001. (E) Venn diagram depicting the number of genes co-bound by THAP1, YY1 and HCFC1 (ChIPseq K652 cells, ENCODE). (F) Genome browser track showing CHIP-Seq signals (peaks) for THAP1, YY1, HCFC1 and H3K9ac (ENCODE datasets in K652 cells) at *Ech1, Ceudc2, Prepl and Gapdh* loci. The top panel shows idiogram and the exon-intron structure for all the loci of interest. (G-H) Quantitative ChIP (qChIP) demonstrating the binding of YY1, and H3K9ac at the promoter region of *Ech1, Cuedc2, Prepl and Gapdh* loci in NSC. Binding, represented as % input (y-axis), is demonstrated for (G) YY1 and (H) H3k9ac for chromatin isolated from *Thap1*^*+/+*^, *Thap1*^*F81L/-*^ and *Thap1*^*-*/-^ NSC cells for the promoter region of *Ech1, Cuedc2, Prepl and Gapdh* loci (x-axis); One-way ANOVA for YY1 qChIP - *Ech1* - F(2,3) = 1877, p<0.0001; *Cuedc2* - F_(2,3)_ = 624.9, p<0.0001 and *Gapdh* - F_(2,3)_ = 12.48, p=0.0351; Dunnett’s multiple comparisons test: adjusted p value < 0.0001. One-way ANOVA for H3K9ac qChIP - *Ech1* - F_(2,3)_ = 35.78, p = 0.0081; *Cuedc2* - F_(2,3)_ = 35.32, p = 0.0082 and *Gapdh* - F_(2,3)_ = 11.72, p = 0.038; Dunnett’s multiple comparisons test: adjusted p value < 0.0001.

Given these observations, we tested whether the F81L mutation impairs THAP1 function in NSC, as it does in CNS tissue. We compared the expression of the THAP1 target genes *Ech1, Cuedc2, Prepl* and *Gapdh in Thap1*^*F81L/-*^, *Thap1*^*+/+*^, and *Thap1*^*-/-*^ NSCs. Compared to controls, *Thap1*^*F81L/-*^ NSCs exhibited significantly lower expression of all THAP1 target genes assessed (> 2 fold decrease in *Thap1*^*F81L/-*^ for *Ech1, Prepl* and *Cuedc2;* ANOVA: p < 0.0001; Figure 3D). These findings are consistent findings those in DYT6 cerebral cortex (Figure 1E) and indicate that the F81L mutation impairs the transcriptional function of THAP1 via a mechanism distinct from DNA binding.

### DYT6 mutations disrupt the THAP1-mediated transcriptional complex at target loci

Significantly reduced transcription of core THAP1 target genes despite normal *Thap1*^*F81L*^ DNA binding at their promoters suggests that the F81L mutation impairs the binding of THAP1 transcriptional partners. We used *in silico* analyses to identify the candidate THAP1 transcriptional partners. We utilized ChromNet, which uses a conditional-dependence network among regulatory factors from ENCODE ChIP-seq data sets to identify closely related datasets (17). A network of YY1, HCFC1 and H3K9ac were the strongest predicted THAP1 interactors, based on shared genome location (Figure S2). These analyses demonstrate that 80.9% and 82.7% of the promoter regions of all THAP1-target genes are co-bound with YY1 and HCFC1, with 71.3% of genes co-bound with both YY1 and HCFC1 (venn diagram, Figure 3E). Consistent with our findings these datasets predicted strong enrichment of YY1, HCFC1 and H3K9ac specifically for all our core THAP1 target genes (*Ech1, Cuedc2, Dpagt1* and *Prepl)* (Figure 3F).

Several lines of evidence led us to focus on YY1 as potentially impacted by the F81L mutation. YY1 is a transcription factor known to interact with THAP1 (4), has a known role in OPC differentiation (18), and is implicated in dystonia (19-22). We also examined THAP1-dependent changes in H3K9ac, an epigenetic marker of active transcription that is highly associated with THAP1-bound chromatin. qChIP in NSC confirmed YY1 binding and H3K9ac enrichment at the promoter regions of *Ech1, Cuedc2* and *Prepl* in NSC (Figure 3G-H). A signal for H3K9ac, but not YY1, was detected at the *Gapdh* locus (Figure 3G-H), validating the *in silico* analyses. Strikingly, there was a significant loss of YY1 binding (Figure 3G) from the promoters of *Ech1, Cuedc2* and *Prepl* in *Thap1*^*F81L/-*^ *and Thap1*^*-/-*^ NSC (> 75% decrease in *Thap1*^*F81L/-*^ and > 90% decrease *Thap1*^*-/-*^ for *Ech1, Cuedc2* and *Prepl* relative to *Thap1*^*+/+*^; ANOVA; t p<0.001), that correlated with decreased transcription from these loci (Figure 3A). As with YY1, we observed a significant reduction of H3K9ac enrichment (Figure 3H) in DYT6 *Thap1*^*F81L/-*^ NSC (*>* 50% decrease in *Thap1*^*F81L/-*^ and > 75 % loss in *Thap1*^*-/-*^ NSC relative to *Thap1*^*+/+*^ for *Ech1 and Cuedc2* ; ANOVA; p<0.001; for *Prepl* ; ANOVA; p<0.05). No differences were observed for enrichment of H3K9ac at the *Gapdh* loci in *Thap1*^*F81L/-*^ *or Thap1*^*-/-*^ NSC, demonstrating specificity at THAP1 target genes. These findings suggest strongly that the F81L mutation acts by impairing the ability of THAP1 to normally assemble an active transcriptional complex that includes YY1.

## Discussion

Our studies establish that the F81L mutation impairs THAP1 transcriptional function and causes *in vivo* defects in myelination similar to those observed in CNS conditional THAP1 knockout mice. Our studies demonstrate that the F81L missense mutation prevents THAP1 from organizing a YY1-containing co-activator complex but does not significantly alter DNA binding.

Prior structural studies of the THAP domain predict that the F81 residue does not participate in DNA binding (6, 15). Our qChIP data are consistent with that prediction, demonstrating no change in binding of endogenous levels of F81L mutant THAP1 at its target loci. This behavior appears distinct from that caused by the C54Y missense mutation, which mutates the Zn binding C2CH motif of the THAP domain. In contrast to *Thap1*^*F81L/-*^ mutants, *Thap1*^*C54Y/-*^ mutant mice exhibit embryonic lethality (23), because of a marked impairment of DNA binding (10, 14).

The significant reduction in DNA binding of YY1 was paralleled by the loss of H3K9ac histone modification. These observations support the conclusion that F81L acts by impairing the ability of THAP1 to organize an active transcriptional complex. In prior studies, we and others (4, 10) have identified several transcription factors and epigenetic modifiers that are significantly co-enriched with THAP1 on the genome. Most prominent among them are YY1 and HCFC1 (Figure 3E), proteins that interact with THAP1 (4, 24, 25) and co-regulate THAP1-bound genes (4, 26).

Multiple aspects of YY1 function are consistent with a key role in DYT6 pathogenesis. There is extensive co-binding of YY1 and THAP1 at the genomic level (4), and these proteins play a co-regulatory role in transcription (4). Both proteins also participate in CNS myelination via cell autonomous effects in the oligodendrocyte lineage (18). Several recent studies report YY1 loss-of-function mutations in dystonia patients (19-22). Considered together, these findings suggest that these two transcription factors operate in a shared pathway impacting CNS myelination and motor function. Prior work identifying the relationship between THAP1 and YY1 was performed in THAP1 null mice (4). Our new findings extend that work, linking abnormalities of YY1 binding to a pathogenic, disease causing DYT6 mutation. We expect that the binding of additional THAP1 co-factors (e.g., HCFC1) are also impaired by THAP1^F81L^. Indeed, prior studies demonstrate that when expressed in SH-SY5Y, several DYT6 mutants (p.N136S, p.N136K, p.Y137C) exhibit functional deficits in recruiting HCFC1 (25).

Identification of the F81L-THAP1 as a hypomorphic protein has important implications for understanding the dominant inheritance of the disease. As THAP1 functions as a dimer (27-29), our findings raise the possibility that THAP1^F81L^ exerts a dominant-negative effect on the wild type protein. Exploring whether F81L and wild type THAP1 interact, and the transcriptional impact of such a heterodimer are important future considerations that will further advance our understanding of the molecular pathogenesis of DYT6 dystonia.

## MATERIALS & METHODS

### Generation and maintenance of mice

Animal research was conducted in accordance with the NIH laboratory animal care guidelines and with the Institutional Animal Care and Use Committee (IACUC) at the University of Michigan. Generation, characterization and genotyping of knock-in mice containing a floxed *Thap1*^*F81L*^ allele in exon 2 have been previously described (4). Nestin-*Cre*+ was purchased from Jackson Laboratory (Stock # 003771). The breeding strategy used to derive all primary NSC cells and conditional null mice was as follows: *Thap1*^*F81L/–*^, *Nestin-Cre*^*+*^ was crossed with *Thap1*^*F81L/+*^ or *Thap1*^*F81L/+*^ to produce the following genotypes: *Thap1*^*+/+*^; *Thap1*^*+/-*^; *Thap1*^*F81L/–*^; *Thap1*^*F81L/+;*^ *Nestin-Cre*^*+*^; *Thap1*^*F81L/–;*^ *Nestin-Cre*^*+*^(N-CKO); *Thap1*^*F81L/–*^, *Nestin-Cre*^*+*^ (N-CKO). Age and sex-matched littermate mice were used for all experiments. Primers used for genotyping in this study (*Thap1* and *Cre*) are listed in Table S2.

### RNA extraction and qRT-PCR

Total RNA extraction from NSC cultures for qRT-PCR and RNAseq analysis was done using NucleoSpin® RNA (Takara) and Trizol (Thermofisher) for mouse cerebral cortex as per manufacturers instructions. cDNA synthesis from total RNA was done using MMLV Reverse Transcriptase (Takara) as per manufacturers instructions. Quantitative real time PCR (qRT– PCR) was performed with the StepOnePlus System (ABI) and 2x SYBR Power Mix (ABI). Primers used for gene expression analyses in this study are listed in Table S2.

### Electron microscopy

EM was performed as previously described (4). P21 mice were anesthetized and perfused with 3% paraformaldehyde / 2.5% glutaraldehyde in 0.1M phosphate buffer. Brains were dissected and postfixed at 4°C overnight. Tissues were dissected, processed and sectioned at Emory University EM core facility. EM image acquisition was done at the University of Michigan, MIL core services using JEOL JSM 1400.

### Automated analysis of g-ratio and axon caliber

Axon caliber (average of the major and minor axes of the axonal perimeter) and g-ratio (ratio of the inner axonal diameter to total (including myelin) outer diameter) were calculated using an automated approach as follows: for each EM image acquired at 20,000x magnification, the inner and outer perimeters of myelinated axons were manually traced and saved as binary masks using ImageJ. The g-ratio and axon caliber of each myelinated axon were calculated using CellProfiler 3.1.9 (30). For each axon, the major axis, minor axis, and length of inner and outer perimeter were measured using the manually traced binary masks. G-ratio was calculated by dividing the inner perimeter by the outer perimeter, and axon caliber was calculated by taking the average of the major and minor axes of the inner perimeter.

### Derivation of NSC cells

NSC were isolated from the sub ventricular zone (SVZ) of P7 mouse pups corresponding to *Thap1*^*+/+*^; *Thap1*^*F81L/–*^ or *Thap1*^*F81L/–;*^ *Nestin-Cre*^*+*^(N-CKO) genotypes as previously described (4, 31) to be propagated and maintained as clonal lines. SVZ derived neurospheres were propagated in NSC growth media (Neurobasal media supplemented with 1x B27, 1x Antibiotic-Antimycotic, 1x Glutamax, 20 ng/µl Fgf2 and 20 ng/µl Egf2) in low attachment T-25 flasks for the first two passages. NSC clonal lines were further expanded as a monolayer on laminin coated dishes in NSC expansion media (DMEM/F12 media supplemented with 1x N2, 1x AA, 1x Glutamax, 20 ng/µl Fgf2 and 20 ng/µl Egf2).

### Quantitative chromatin immunoprecipitation (qCHIP)

qCHIP was performed as previously described (32). Sheared chromatin (sonicated to 200–500 bp) from 2 × 10^6^ mouse NSCs was incubated with 4μg of Goat THAP1 (sc-98174, Santa Cruz), 4μg of Rabbit YY1 (sc-98174, Santa Cruz), 2μg of Rabbit H3K9ac (sc-98174, Activemotif) or 4μg of normalized Goat IgG (Santa Cruz), or normalized 4μg of Rabbit IgG (Santa Cruz) using Dynabeads (ThermoFisher Scientific). After washing, elution and cross-link reversal, DNA from each ChIP sample and the corresponding input sample was purified (PCR Cleanup, Takara) and analyzed further using qPCR. Each ChIP sample and a range of dilutions of the corresponding input sample (0.01 – 2% input) were quantitatively analyzed with gene-specific primers using the StepOnePlus System (ABI) and SYBR qPCR Powermix (ABI). Primers used for qChIP analyses in this study are listed in Table S2.

### Statistics

All data are reported as mean ± SEM. All statistical tests reported (Student’s t-tests, One-way ANOVAs) were performed using Graphpad Prism software (V9).

## Data & software availability

The GEO accession number for CHIP-seq data used in manuscript for THAP1 is GSM803408, YY1 is GSM803446 and H3K9ac is GSM788082. The GEO accession number for gene expression data for *Thap1* cKO CNS tissue is GSE97372 and *Thap1* cKO oligodendrocyte lineage is GSE161556. The code utilized for automated analysis of g-ratio and axon caliber using CellProfiler 3.1.9 is available at https://github.com/suminkim/Analysis_CellProfiler.

## Acknowledgements

We thank Haoran Huang and Jack Koulos for technical assistance. We thank Dr. Vikram Shakkottai and Dr. Roman Giger for assistance with lab equipment and microscope. We thank Cathy Collins for critical reading of manuscript. We thank the staff of University of Michigan’s Core Facilities (DNA Sequencing Core, Microscopy and Image Analysis Laboratory and Unit of Laboratory Animal Medicine) and Hong Yi (EM Core, Emory University). This research was supported in part by the following grants: to WTD (1R01NS109227 NINDS).

## Abbreviations

THAP1: Thanatos-associated [THAP] domain containing associated protein 1
OL: oligodendrocyte
OPC: oligodendrocyte progenitor cells
ENCODE: The Encyclopedia of DNA Elements
ChIP: Chromatin immunoprecipitation

