## Supplemental Figures for "A pathogenic DYT-THAP1 dystonia mutation causes hypomyelination and loss of YY1 binding"

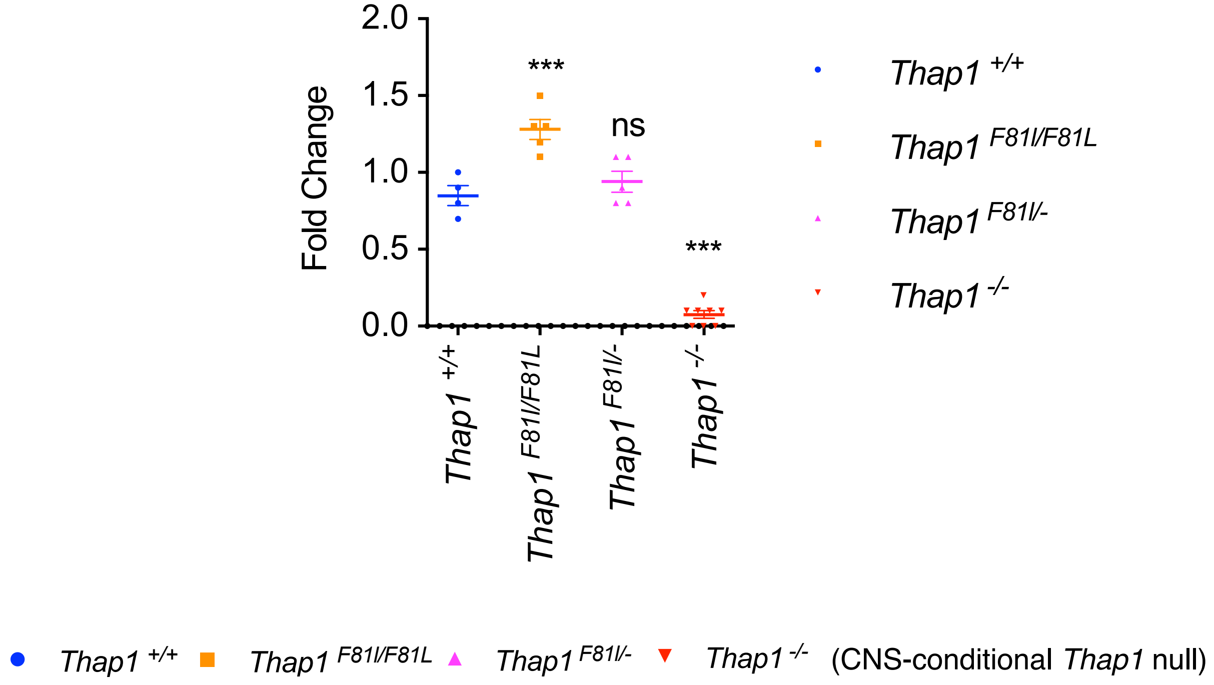


**Figure S1:** No reduction in the mRNA expression of *Thap1* in *Thap1*$^{F81L/F81L}$ and *Thap1*$^{F81L/-}$ CNS (cerebral cortex) relative to control (*Thap1*$^{+/+}$) as measured by qRT-PCR. mRNA expression for each individual gene was normalized to *Rpl19* expression and represented in the bar graph (mean ± SEM) as fold change (y-axis) for all genotypes (x-axis) with respect to *Thap1*$^{+/+}$. One-way ANOVA for *Thap1* = $F_{(3,18)}$ = 117.3, $p<0.0001$; Dunnett's multiple comparisons test: adjusted p value = 0.0002 for *Thap1*$^{+/+}$ vs. *Thap1*$^{F81L/F81L}$ ; adjusted p value = 0.54 for *Thap1*$^{+/+}$ vs. *Thap1*$^{F81L/-}$; adjusted p value < 0.0001 for *Thap1*$^{+/+}$ vs. *Thap1*$^{-/-}$

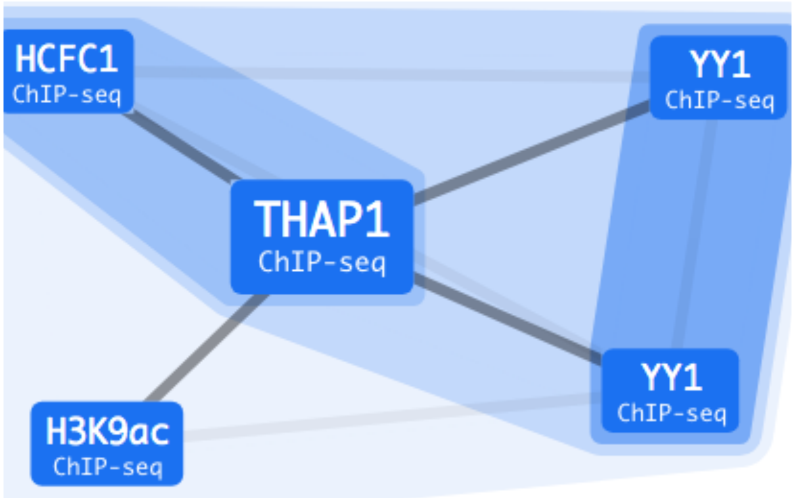


**Figure S2:** Screen capture from the web interface of ChromNet (http://chromnet.cs.washington.edu), used to identify the ChIPseq datasets interacting with that of THAP1 with a correlation greater than 0.9. THAP1 ChIPseq dataset used for this analysis was previously deposited in ENCODE database and derived from K652 cells. THAP1-interacting datasets (YY1, HCFC1 and H3K9ac) as identified by ChomNet were all limited to K652 cells and previously deposited in ENCODE. Details and accession numbers for the datasets used have been provided in the manuscript. The edge threshold was set at 0.4 and captured the three strongest edges (datasets) connected to THAP1.
